# Exercise preconditioning confers skeletal muscle myometaplasticity

**DOI:** 10.64898/2026.09.20.753020

**Authors:** Toby L. Chambers, Ana Regina Cabrera, Ronald G. Jones, Pieter J. Koopmans, Francielly Morena, Nathan Serrano, Aram Parhizkar, Calvin S. Peterson, Abbey L. Stokes, Zain B. Malik, Konstantinos Papanikolaou, Ferdinand von Walden, Davis A. Englund, Cory M. Dungan, Christopher E. Nelson, Yuan Wen, Kevin A. Murach

## Abstract

Previously exercise trained skeletal muscle is more growth-responsive to retraining. Using a murine training-detraining-retraining approach and multi-omics, we find that *Pecam1* gene expression is lower, but capillarization is appreciably higher in previously trained (preconditioned) relative to naïve trained control muscle. Greater capillarity could be permissive for accelerated hypertrophic adaptation. Exercise preconditioned myonuclei feature differential promoter CpG regulation in genes related to *Wnt* signaling. These epigenetic alterations with retraining align with our prior observations of methylation changes within the same pathway after a longer period of chronic training, suggesting a more rapid response due to preconditioning. Methylome-transcriptome integration and single myonucleus RNA-sequencing expose the polyamine metabolism enzyme *Smox* as a target that relates to heightened hypertrophic adaptability with retraining*. Smox* induction is sufficient to cause hypertrophy in aligned myotubes cultured on a stiffness-tuned substrate along with a growth-supportive transcriptional program. Integration of our multi-omics data suggests that *Smox* regulates repression of *Ddit4/Redd1* (an inhibitor of mTORC1 signaling) after retraining. *Smox* may govern a favorable muscle fiber growth environment in previously trained muscle by sensitizing anabolic potential through polyamine metabolism. A lower adaptive threshold mediated by *Smox* could contribute to myometaplasticity, or a change to how subsequent muscle adaptations are made.

## 1. INTRODUCTION

In neuroscience, a priming stimulus can shift the threshold for subsequent long term neural potentiation or depression, known as neural “metaplasticity” [1; 2]. Skeletal muscle fibers are post-synaptic, highly differentiated, and excitable cells that share this same potentiation quality as neural cells. The triggers and mechanisms driving cellular muscle metaplasticity, or “myometaplasticity”, are beginning to be explored but are not well-defined. Metaplasticity is distinct from memory. Memory is the retained adaptation, whereas metaplasticity is a change in how the next adaptation is made. Emerging evidence suggests that prior exercise training primes skeletal muscle for enhanced adaptability in the future [3–7]. Priming of the molecular machinery via several possible mechanisms may explain more rapid and/or augmented hypertrophic adaptations observed in exercise retrained or “preconditioned” skeletal muscle compared to naïve trained skeletal muscle [3; 4; 6]. Understanding the mechanisms for how skeletal muscle is naturally sensitized by prior exercise training has implications for athletic performance following cessation from training (e.g. due to injury), as well as for combatting conditions of compromised muscle health.

Our previous work identified DNA methylation signatures specifically in muscle fiber nuclei (myonuclei) from previously trained mice after detraining [6]. This epigenetic signature suggests a “muscle memory” of prior exercise that persists following the return to a pretrained cellular phenotype [6; 8], consistent with what is observed in human skeletal muscle tissue after detraining [3–5; 9]. The myonuclear epigenetic muscle memory we observed preceded an enhanced whole muscle and cellular hypertrophic response to retraining [6], in alignment with similar observations after exercise retraining in humans [4; 5; 10]. In the current investigation, we profiled exercise primed skeletal muscles from our murine study after retraining and compared them to age-matched exercise-naïve controls that underwent volume-matched exercise training [6]. There are several possible mechanisms that could explain the molecular bases of muscle memory (e.g, persistent DNA methylation, histone and chromatin modifications, miRNA expression, long-term proteome changes, etc.) [3; 4; 11-14] that then contribute to myometaplasticity. Here, we used myonuclear DNA methylation profiling to be consistent with our prior work [6] but combined it with a multi-pronged analysis focused on myonuclei and their transcriptional output in the retrained state. We hypothesized that this muscle fiber-centric exploratory omic approach would reveal new molecular regulators of hypertrophic sensitivity that relate to myonuclear DNA methylation status or is independent from it.

Several methodological advances in this study permit detailed investigation of how exercise-sensitized skeletal muscle leads to enhanced hypertrophic responsiveness: 1) a translatable method of high-volume endurance and resistance exercise training in mice – progressive weighted wheel running, or PoWeR [8; 11; 15; 16] – that enables training-detraining-retraining studies in mice, 2) a recombination-independent doxycycline-inducible method (HSA-rtTA-TetO-GFP, or HSA-GFP) for labelling resident myonuclei at the onset of training [6; 11; 17-19], 3) low-input myonucleus-specific DNA methylation analysis and single myonucleus RNA sequencing (smnRNA-seq) on an enriched myonuclear population, 4) a multi-omic integration approach that links DNA methylation to gene expression that was previously validated against functional cellular outcomes [20; 21] and 5) an *in vitro* approach for generating mature and aligned myotubes on a stiffness-tuned substrate [22] that are then manipulated using myotropic adeno-associated viruses (myoAAV) [23] for candidate gene target validation. Altogether, we define the molecular characteristics of retrained skeletal muscle in granular detail and propose that exercise preconditioning may maintain a lower threshold for anabolic potential rather than by amplifying the growth stimulus. Within this framework, enhanced capillarity, Spermine oxidase (*Smox*)- dependent polyamine metabolism, and *Ddit4/Redd1* suppression may contribute to superior hypertrophic responsiveness and myometaplasticity. Our findings could have therapeutic implications for skeletal muscle pathologies (e.g. rehabilitation, injury, aging, etc.) in clinical populations and/or improve muscle performance in the context of athletic competition.

## 2. RESULTS

### 2.1 Differential myonuclear DNA methylation of Wnt genes is associated with enhanced hypertrophic responsiveness upon retraining

An overview of the experimental design is presented in Figure 1A. In brief, per our prior study [6], adult male HSA-GFP mice performed voluntary PoWeR for 8 weeks then detrained for 12 weeks, before a retraining period of 4 weeks (DTRT). Training-naïve controls (CON) were age-matched mice (not trained previously) and performed voluntary PoWeR for 4 weeks. Mice were ∼9 months old at the start of the 4 weeks of PoWeR training and ∼10 months old at study completion. Mice were treated with low-dose doxycycline in drinking water (0.5 mg/mL with 2% sucrose) to fluorescently label resident myonuclei with GFP for 1 week prior to the 4-week PoWeR training program for both DTRT and training naïve CON animals (Figure 1A). The timing of myonuclear labeling captures “resident” myonuclei and ensures the exclusion of muscle stem cells (satellite cells) accrued during the PoWeR training period. Muscle was collected 48 hours after the final exercise bout in an overnight fasted state. Exercise preconditioning (DTRT) resulted in differential methylation (DM) in ∼7.2% of measured myonuclear promoter CpG sites (defined as CpG sites located within 10% of corresponding gene length around the transcriptional start site using the exon 1 start codon as a reference position) assessed by reduced representation bisulfite sequencing (RRBS) versus myonuclei from naïve trained mice. Of the DM CpG sites in predicted promoter regions in exercise preconditioned muscle, 17257 and 38623 were hypo- and hypermethylated, respectively, compared to naïve (*q*<0.05, Supplemental Table S1). These 55880 DM CpG sites where present across 8829 genes.

**Figure 1.**
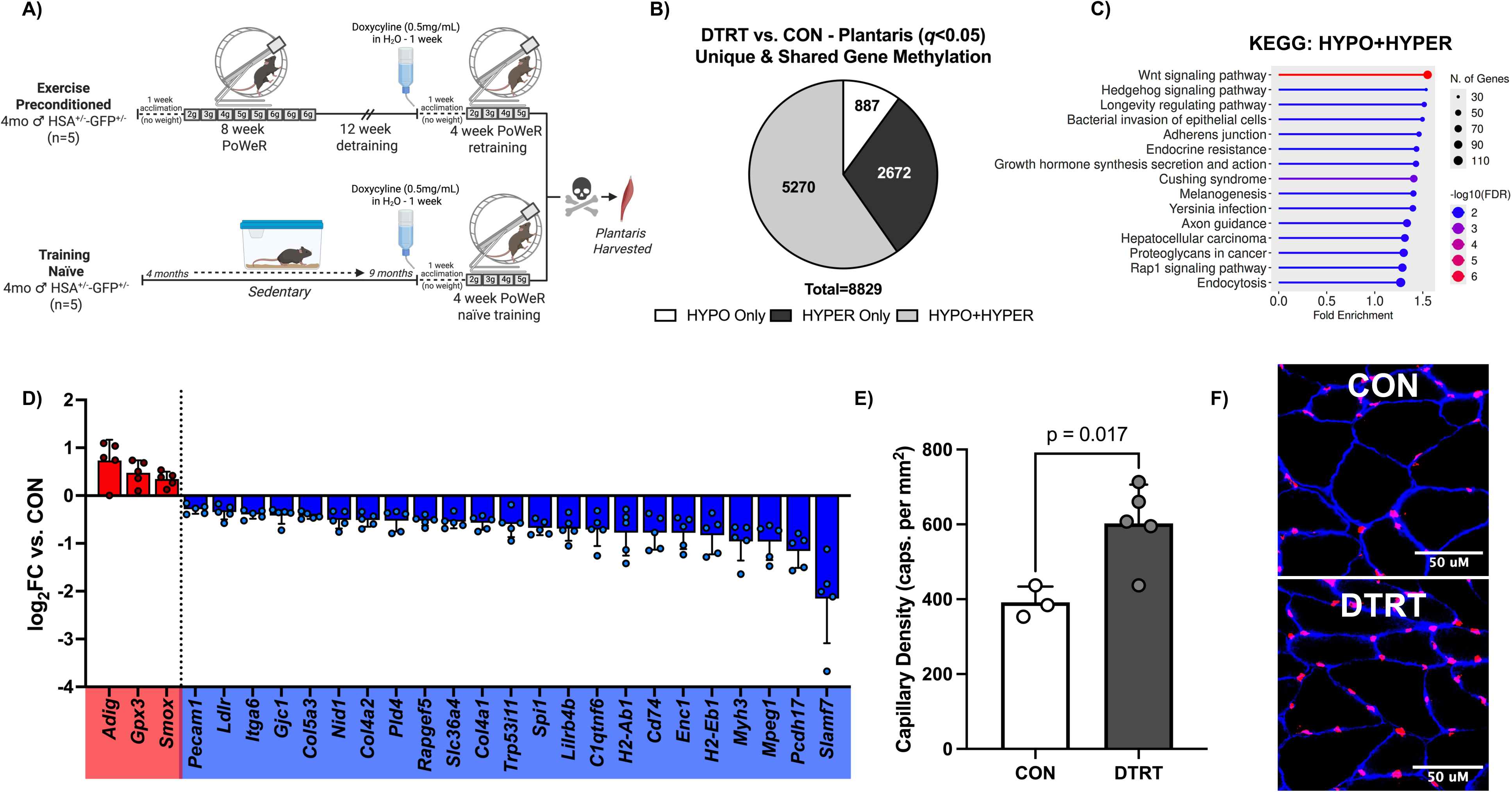
Study overview and epigenetic regulation of the whole muscle transcriptome with exercise preconditioning. (A) Schematic overview of experimental design. (B) Number of genes with unique (HYPO or HYPER only) and simultaneous (HYPO+HYPER) DNA methylation in promoter regions. (C) KEGG pathway enrichment of genes with simultaneous HYPO+HYPER DNA methylation in their promoter regions. (D) Plot of log_2_FCs of up- and downregulated BETA targets inferred to be epigenetically regulated. (E) Plantaris muscle capillary density and (F) representative IHC images (pink=PECAM1+/capillary; blue=dystrophin).

To assess genes with unique (hypo- or hypermethylation only) or simultaneous (hypo+hyper) DNA methylation in promoter regions, we collapsed the data into three subcategories: 1) genes with CpG sites in the promoter region that were hypomethylated only, 2) hypermethylated only, and 3) mixed methylation (hypo+hyper). Of the 8829 DM genes, 887 and 2672 had unique hypo- and hypermethylation DNA methylation at their CpG promoter regions, respectively. The remaining 5270 genes had both hypo- and hypermethylated CpGs in their promoter regions (Figure 1B, *q*<0.05). KEGG analysis yielded no significant enrichment for genes with unique (hypo or hyper only) DNA methylation. However, KEGG analysis of genes with simultaneous DNA methylation (hypo+hyper) yielded enrichment of *Wnt* and Hedgehog signaling, as well as “longevity regulation”, adherens junction, endocrine resistance, axon guidance, and endocytosis pathways (Figure 1C). Our findings reinforce the complex nature of epigenetic regulation at promoter region CpG sites during retraining. These findings also suggest that enhanced hypertrophy upon retraining (∼10% greater muscle mass versus CON [6]) may be regulated, in part, by differential myonuclear promoter DNA methylation and potentially fine-tuned gene expression, particularly in the *Wnt* signaling pathway.

### 2.2 Methylome-transcriptome integration identifies regulation of capillarization as a characteristic of enhanced hypertrophy in retrained muscle

The relationship between gene expression and DNA methylation is at times seemingly paradoxical, leading to an unclear understanding of how DNA methylation controls gene expression [24–26]. For example, promoter hypomethylation does not always correlate to an increase in the expression of a given gene, as would be expected [27–29]. Since the role of DNA methylation on transcriptional regulation is incompletely defined, we employed a holistic computational approach called Binding and Expression Target Analysis (BETA) that considers methylation patterns (hypo and hyper) around a gene as well as proximity to the transcription start site to infer transcriptional regulation [20; 21; 30]. During muscle adaptation, we report that this approach infers epigenetic regulation of transcripts that corresponds to measurable muscle function outcomes [20], reinforcing the veracity of this method.

BETA analysis inferred myonuclear epigenetic regulation of bulk gene expression in preconditioned compared to naïve trained skeletal muscle (Figure 1D, adj. *p<*0.05). Upregulated epigenetically-controlled genes in preconditioned muscle were related to polyamine metabolism (*Smox*), lipid metabolism (*Adig*), and cellular redox (*Gpx3*) (Figure 1D, adj. *p<0.05*). Downregulated genes related to the extracellular matrix (*Col4a1, Col5a3, Itga6, Nid1*), immunity (*Slamf7, Mpeg1, H2-Eb1, H2-Ab1, Cd74, Lilrb4b, Spi1, Pld4),* and capillarity (*Pecam1*) were inferred to be epigenetically regulated in exercise preconditioned compared to naïve muscle after 4 weeks of training. While PECAM1 (CD31) labels blood vessels and is highly enriched in endothelial cells, it can be expressed sporadically in myonuclei as well [31].

Lower *Pecam1* expression in the bulk RNA-seq data set did not correspond to reduced capillarity in preconditioned muscle. Perhaps paradoxically, plantaris capillary density was 53.6% higher in preconditioned than naïve muscle (602 vs 392 capillaries/mm², Figure 1E-F, *p*<0.05). Capillary density with detraining and retraining is therefore decoupled from steady-state *Pecam1* transcript abundance. Whether this reflects endothelial persistence from the initial training period, angiogenic responsiveness to retraining, or both, the resulting vascular bed may be permissive for growth without requiring sustained *Pecam1* induction. Several studies in humans implicate capillary adaptations to hypertrophic responsiveness with exercise [32–35]. In humans, capillarization may be maintained after ∼12 weeks (84 days) following the cessation of endurance training in the gastrocnemius and vastus lateralis, suggesting long-term capillary maintenance during periods of detraining is possible [36]. The maintenance of capillaries with detraining is not a universal finding, however [37; 38].

### 2.3 Single myonucleus RNA sequencing (smnRNA-seq) exposes the muscle fiber-specific transcriptional landscape of enhanced hypertrophic adaptability with exercise preconditioning

To further explore the potential cell-autonomous effects of exercise preconditioning on skeletal muscle adaptation during retraining, we performed smnRNA-seq on resident myonuclei (i.e. myonuclei labelled fluorescently prior to the 4-week PoWeR training period) in exercise preconditioned and naïve trained animals. The benefit of smnRNA-seq is nucleus type-specific resolution as well as enhanced fidelity compared to bulk RNA-seq that can disguise changes in myonuclei. Plantaris muscle resident myonuclei were isolated using fluorescence-activated nuclear sorting (FANS), as we have described previously [6; 39–41]. Replicates (n=2 per group) were generated by pooling isolated myonuclei from 2-3 plantaris samples from each group and used for the exploratory smnRNA-seq experiment. From the pooled samples (n=2 each for DTRT and CON; total=4), 13653 individual resident myonuclei were isolated, sequenced, categorized, and mapped based on their expression profile.

The integrated UMAP of resident plantaris myonuclei resolved six major myonuclear populations: spindle, Type IIa, Type IIx, Type IIb, neuromuscular junction (NMJ), and myotendinous junction (MTJ) (Figure 2A). Myonuclear subtypes were annotated using curated panels that included myosin heavy chain isoform transcripts as dominant markers alongside supporting cluster-specific genes, with muscle spindle myonuclei accordingly most enriched for *Myh7b, Myh6,* and *Myl2,* Type IIa for *Myh2*, Type IIx for *Myh1*, Type IIb for *Myh4*, NMJ for *Col19a1* and *Colq,* and MTJ for *Col22a1*. Notably, Type I myonuclei were not identified in the dataset, consistent with the well-established near-complete absence of MyHC I muscle fibers in rodent plantaris [42]. Muscle spindle-, NMJ-, and MTJ-associated myonuclei clusters were small in the current investigation and yielded minimal DEGs between preconditioned and naïve skeletal muscle (Supplemental Table S2). Our primary findings are therefore focused on the coarse myonuclear comparisons between exercise preconditioned and naïve. This is a composite of all assessed myonuclei clusters: 95% were derived from Type II-associated myonuclei (Type IIa: 24.1%; Type IIx: 43.9%, Type IIb: 28.9%; Total Type II: 96.9%). Complete lists of DEGs and enrichment analyses for each myonuclei cluster are available in the supplemental information (Supplemental Table S2).

**Figure 2.**
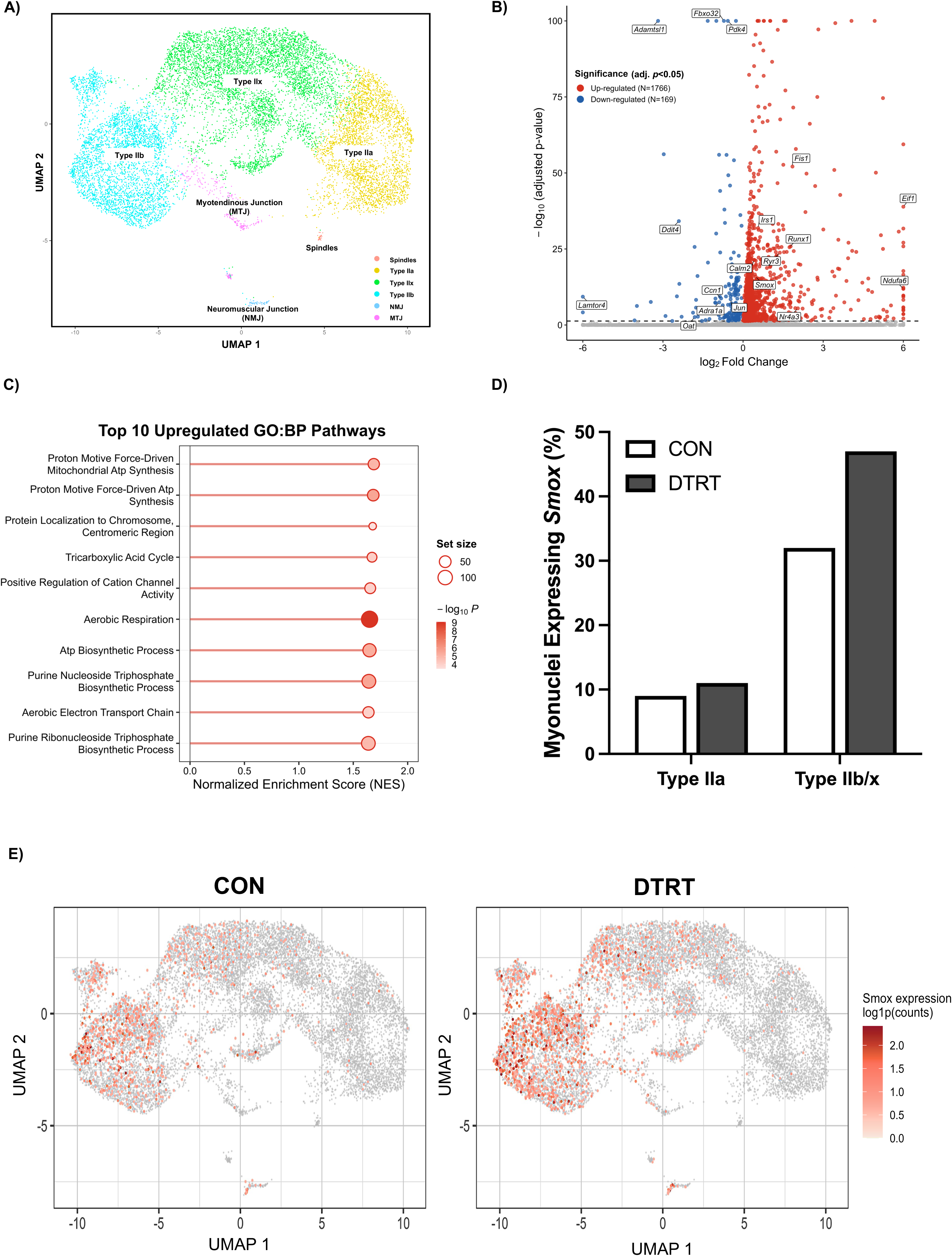
Single myonucleus RNA-sequencing (smnRNA-seq) analysis between naïve trained and exercise preconditioned resident myonuclei. (A) Integrated UMAP of all experimental conditions, labeled and colored by myonuclear cluster. Spindles = muscle spindles. Myonuc_IIA = Type IIA. Myonuc_IIB = Type IIb. Myonuc_IIx = Type IIx. Myonuc_. (B) Volcano plot of all differentially expressed genes in the coarse myonuclear smnRNA-seq dataset (adj. *p*<0.05). (C) Top 10 upregulated GO biological processes based on coarse smnRNAseq myonuclear cluster. (D) Normalized expression and percentage of myonuclei expressing *Smox* in Type II smnRNAseq clusters. (E) Split UMAP of groups specific *Smox* expression in individual resident myonuclei. MTJ, myotendinous junction; NMJ, neuromuscular junction.

Coarse per myonucleaus differential expression (composite smnRNA-seq expression of all myonuclei) revealed appreciable remodeling of the myonuclear transcriptome: 1766 genes were upregulated and 169 were downregulated (Figure 2B, adj. *p*<0.05). In preconditioned myonuclei, eukaryotic translation initiation factor 1 (*Eif1)* was the most upregulated gene compared to naïve myonuclei (adj. *p*=1.27E-39, log_2_FC=10.14). EIF1 is a universally conserved protein critical for the initiation of translation by binding to 40S ribosomal subunits to facilitate accessibility to the pre-initiation complex [43; 44]. Gene ontology (GO) analysis of upregulated genes in the coarse smnRNA-seq revealed enrichment of proton motive force-driven mitochondrial ATP synthesis, protein localization to chromosome, TCA cycle, positive regulation of cation channel activity, aerobic respiration/electron transport chain, and purine metabolism biological processes in preconditioned myonuclei (Figure 2C). These data suggest that transcriptomic remodeling of resident myonuclei from previously trained muscle have a signature that may potentiate bioenergetic capabilities through enhanced aerobic metabolism. This signature could be permissive to the exaggerated muscle growth observed in the preconditioned group [6].

In our prior work, we identified several genes in the myonuclear DNA methylome that had retained promoter region epigenetic methylation patterns following 12 weeks of detraining [6]. Given we have myonuclear transcriptome data in the current investigation, we explored how differential gene expression in preconditioned myonuclei could relate to myonuclear epigenetic regulation of these “muscle memory” genes. Three genes with differential promoter region DNA methylation in myonuclei after detraining were upregulated (coarse smnRNA-seq) in preconditioned myonuclei: *Arid4b*, *Myl12b*, and *Eif1a1*. Some evidence suggests ARID4B has a role in anabolism in myogenic cells [45]. Our previous work linked increased MYL12B protein abundance to enhanced *in vivo* skeletal muscle fatigue resistance and total work capacity in ∼24-month-old male C57BL/6N mice [46], implicating this protein in muscle performance. EIF1A1 is involved in translation elongation and, by extension, protein synthesis [47]. Collectively, elevated expression of these genes in myonuclei that relate to myonuclear DNA methylation regulation in previously trained muscle could contribute to expedited re-adaptation.

### 2.5 Smox in preconditioned skeletal muscle is enriched in myonuclei

We identified *Smox* as an enriched gene in our bulk RNA-seq that was inferred to be controlled by myonuclear DNA methylation. *Smox* encodes spermine oxidase, the polyamine enzyme responsible for the biochemical breakdown of spermine to spermidine in all eukaryotic cells [48–50]. Recent work has linked *Smox* and polyamine metabolism to skeletal muscle health in a variety of contexts [48; 51–55]. Since *Smox* is implicated in muscle mass regulation and was identified by our other analyses, we evaluated *Smox* across myonuclear clusters. The number of myonuclei expressing *Smox* across major myonuclear types was proportionally higher in preconditioned versus naïve trained muscle (Figure 2D). *Smox* was elevated in the Type IIb (+43%, adj. *p*=4.42E-16) and Type IIx (+45%, adj. *p*=0.0002) myonuclei compared to naïve (Figure 2E and Supplemental Table S2). Collectively, we identify *Smox* as a candidate gene that could contribute to enhanced muscle mass accrual in exercise preconditioned skeletal muscle.

### 2.5 Smox induction by myoAAV a promotes myotube growth and a growth-oriented transcriptome

To determine whether *Smox* is sufficient to stimulate muscle growth, we used a myoAAV to deliver *Smox* cDNA to mature C2C12 myotubes. Myotubes were differentiated on a stiffness-tuned substrate that allows for long-term culture of mature and highly aligned myotubes to more closely mimic mature myofibers *in vivo* [22]. Induction of *Smox* starting at 10 days post-differentiation increased *Smox* transcript (Figure 3A) and SMOX protein content (Figure 3B-C) compared to control myotubes (*p*<0.05). The ratio of myosin heavy chain (MyHC) to myonuclei was greater with *Smox* myoAAV treatment compared to controls (Figure 3D-E, *p*<0.05). The induction of SMOX therefore causes hypertrophy *in vitro* in mature myotubes.

**Figure 3.**
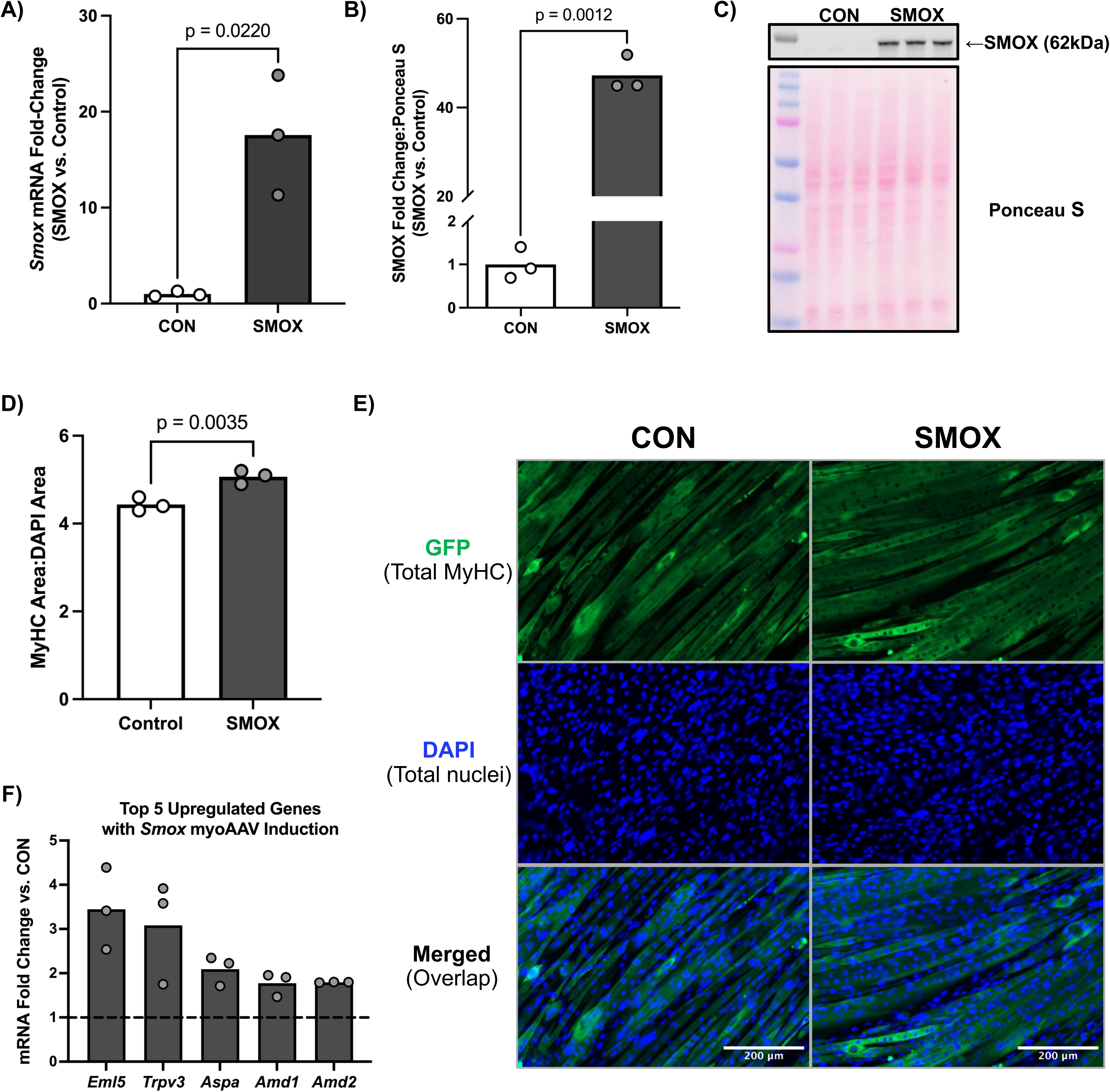
*In vitro Smox* myoAAV induction promotes C2C12 myotube growth and transcriptomic remodeling. (A) *Smox* mRNA expression fold-change in *Smox* myoAAV treated myotubes compared to controls. (B) SMOX protein expression fold-change in *Smox* myoAAV treated myotubes compared to controls and (C) representative Western blot images. (D) Relative myosin heavy chain (MyHC) to DAPI area as a measure of myotube growth with *Smox* myoAAV induction and (E) representative IHC images (green=GFP/Total MyHC, blue=DAPI/Total nuclei). (F) Top 5 most upregulated gene transcripts, by *p*-value, with *Smox* myoAAV induction in mature myotubes.

RNA-sequencing of *Smox*-induced myotubes revealed 741 differentially expressed genes (DEGs) (671 up- and 70 down-regulated, *p*<0.05) compared to scrambled sequence myoAAV (technical triplicates in each condition) (Supplementary Data S3). Adenosylmethionine decarboxylase 1 and 2 (*Amd1/2*) were among the top 5 most upregulated protein-coding genes by *p*-value with *Smox* overexpression (Figure 3F, *p*=0.0002 and *p*=0.0005, respectively). AMD1/2 are enzymes involved in polyamine biosynthesis where *Amd1* is the primary, transcriptionally active protein coding gene and *Amd2* encodes a functional yet less biochemically active enzyme in rodents [53; 56; 57]. Both enzymes catalyze the conversion of S-adenosylmethionine (SAM) to decarboxylated SAM (dcSAM) making it available as a precursor molecule for the production of spermidine and spermine via spermidine and spermine synthases, respectively. Thus, *Smox* induction in mature myotubes appears to be sufficient for hypertrophy, affects polyamine-related gene targets, and provides a potential cell-autonomous mechanistic explanation for how exercise preconditioned skeletal muscle experiences expedited growth upon retraining.

### 2.6 The in vitro Smox overexpression transcriptome & in vivo myonuclear profiles with preconditioning share similarities and reveal Ddit4/Redd1 regulation

We sought to identify common gene programs related to SMOX that could explain enhanced hypertrophic adaptability during retraining. To do this, we cross-referenced DEGs from smnRNA-seq clusters (Coarse, Type IIa, Type IIx, and Type IIb) to the *Smox* myoAAV RNA-seq data (Supplemental Table S4). The number of unique and shared genes between *Smox-*induced myotubes and smnRNA-seq clusters is shown in Figures 4A-D (*p*<0.05). There was overlap for upregulated genes between *in vitro* and *in vivo* experiments for all smnRNA-seq clusters. Of the 671 upregulated genes with *in vitro Smox* overexpression, there was shared expression of 138, 31, 49, and 55 genes between coarse, Type IIa, Type IIx, and Type IIb smnRNA-seq clusters, respectively (Figures 4A-D). Type IIb myonuclei shared the most up-regulated genes with *Smox* myoAAV treatment (Figure 4C), which is the nucleus type with the most significant *Smox* induction in preconditioned muscle (see Figures 2D-E). The literature also suggests that *Smox* is typically higher and more localized in muscle groups with greater distributions of MyHC IIx and IIb myofibers, like the plantaris [58–60]. This fiber type bias for *Smox* may explain the greater overlap of genes between MyHC IIb smnRNA-seq and the *in vitro Smox* induction experiments. *Amd1* was upregulated *in vivo* in MyHC IIb myonuclei, in agreement with the *in vitro Smox-*induced myotubes.

**Figure 4.**
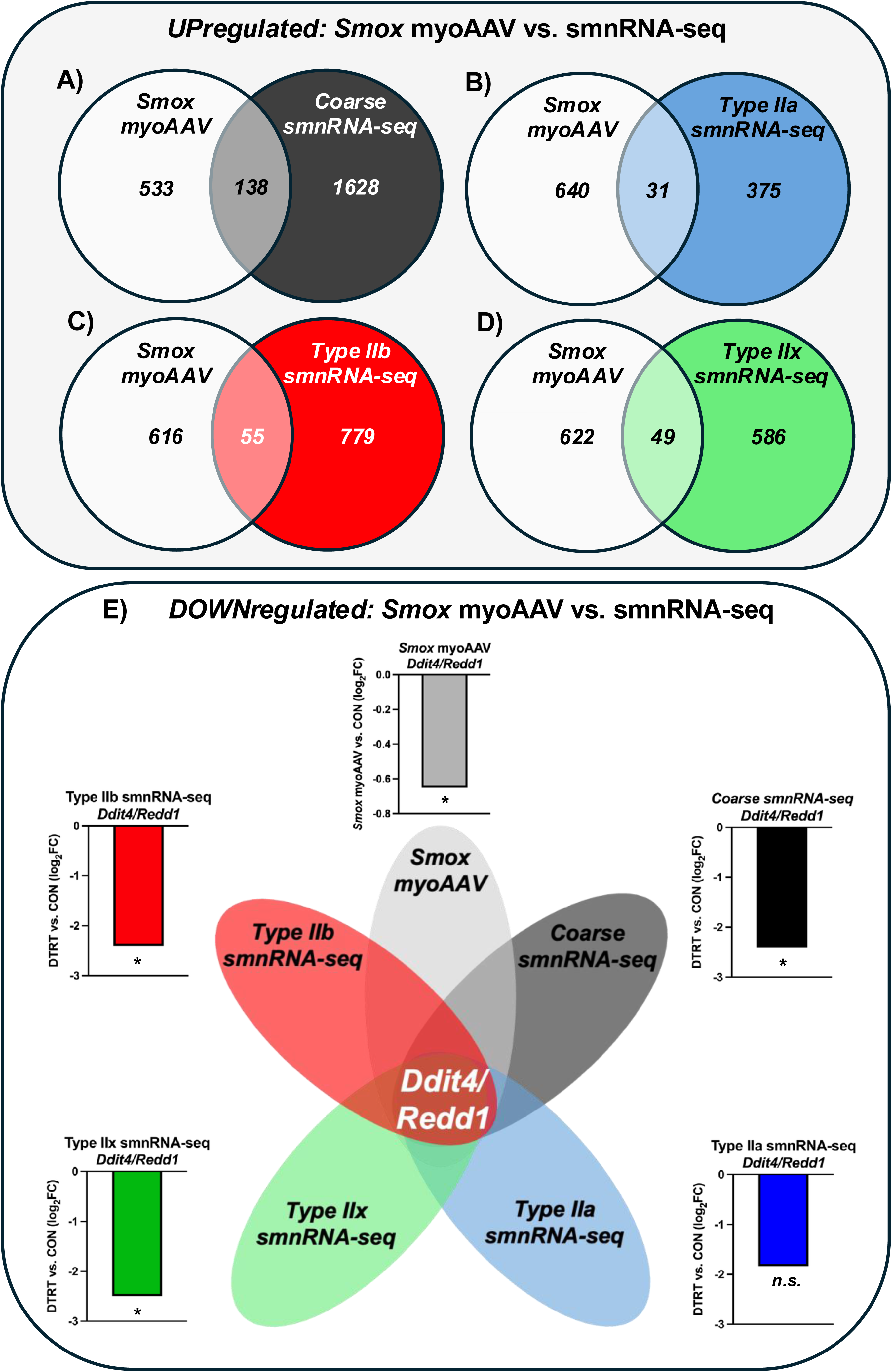
Unique and shared gene expression profiles between the *in vitro Smox* overexpression transcriptome and *in vivo* myonuclear profiles with preconditioning. Overlap of upregulated genes between the *in vitro Smox* overexpression transcriptome and (A) coarse, (B) Type IIa, (C) Type IIb, and (D) Type IIx smnRNA-seq myonuclear clusters. (D) *Ddit4/Redd1* repression between the *in vitro Smox* overexpression transcriptome and smnRNA-seq myonuclear clusters. \**p*- or adj. *p*<0.05; n.s., not significant.

In searching for regulators of muscle mass regulation in our cross-platform analysis, we observed that DNA damage-inducible transcript 4 (*Ddit4/Redd1)* was significantly repressed in preconditioned coarse, Type IIx, and Type IIb smnRNA-seq clusters and in *Smox-*induced myotubes (Figure 4E). *Ddit4/Redd1* encodes a stress-response regulator that inhibits mTORC1 signaling, suppresses protein synthesis, and coincides with muscle wasting in atrophy-associated conditions [61; 62]. Our findings therefore suggest: 1) lower *Ddit4/Redd1* expression in skeletal muscle may contribute to augmented *in vitro* and *in vivo* muscle growth, and 2) increased *Smox* expression indirectly regulates *Ddit4/Redd1* expression, likely through its biochemical function in polyamine metabolism and methionine salvage pathways.

## 3. DISCUSSION

Myonuclei added during training are seemingly lost during detraining in mice and humans [7; 8; 11; 63]. “Muscle memory” may therefore, in part, be regulated by long-lasting epigenetic modifications to resident (non-satellite cell-derived) myonuclei [6; 11]. This epigenetic memory provides a biological blueprint for how muscle re-adaptation can be expedited after a cessation from training in both rodents and humans [4; 6; 7; 9; 40; 64]. Previously, we showed that 8 weeks of PoWeR increased plantaris myofiber size and myonuclear density while shifting fiber type from a more glycolytic to more oxidative metabolic profile in adult male mice [8]. After 12 weeks of detraining, these adaptations were lost and the muscle returned to its initial untrained cellular phenotype; however, there was evidence for a sustained and distinct epigenetic signature of PoWeR training in the myonuclei of detrained muscle relative to untrained [6; 8]. Following the detraining period, previously trained mice underwent 4 weeks of PoWeR retraining where the plantaris muscle hypertrophied to a greater extent compared to training naïve mice (∼10% greater than training naïve) [6]. In the current investigation, the same preconditioned and retrained muscle had differential promoter methylation of genes in the *Wnt* signaling pathway in resident myonuclei. Differential methylation of *Wnt* genes was characteristic of the initial 8-week period [6]. Agreement between our prior 8-week training analysis and current 4-week retraining experiments are supportive of an accelerated training response at the epigenetic level. Also, within this preconditioning-associated molecular rewiring, upregulation of *Smox* occurred simultaneous with repressed *Ddit4/Redd1*. These myofiber-specific molecular alterations may create a muscle growth environment that contributes to enhanced hypertrophy.

Polyamines are small organic polycations with multiple amino groups. They act as essential co-factors that interact with negatively charged DNA, RNA, and proteins to influence DNA methylation and gene expression, stabilize molecular structures, regulate cellular processes, and influence alternative pre-mRNA splicing [49; 65–70]. In mammalian tissue, dysfunctional polyamine metabolism is associated with pathologies related to aging [71–73], sarcopenia [69], cancer [68; 73–75], neurological disorders [76; 77], fibrosis [78], muscular denervation and dystrophy [73; 79–81], and metabolic disease [82]. *In vitro* and *in vivo* studies of skeletal muscle highlight SMOX’s biochemical regulation and role during myogenic cell differentiation [83; 84], muscle development [85], immobilization [54], exercise training [86; 87], and aging [59]. In mouse models of skeletal muscle atrophy and aging, global *Smox* overexpression attenuated muscle mass and contractile losses with concurrent improvements in mitochondrial function compared to control young and old mice [54; 59]. Our findings suggest that *Smox* and polyamine metabolism, in part, regulate the responses in previously trained skeletal muscle that facilitate enhanced growth. This *Smox* regulation appears to be enriched in Type IIb and IIx myonuclei in preconditioned plantaris muscle. Further investigation in the Type I myonuclei of slow-twitch myofibers would help determine if *Smox’s* actions are exclusive to Type II myonuclei and if exercise type (i.e. aerobic vs. resistance) influences these outcomes, as suggested by others [88].

*In vitro* SMOX induction led to higher mRNA abundance of the polyamine enzymes *Amd1* and *Amd2*. This observation inverts the reciprocal relationship in non-muscle cell, where catabolic SMOX activity suppresses AMD1 [89]. Muscle may therefore couple polyamine catabolism and anabolism rather than trading one for the other, sustaining flux through the pathway instead of depleting it. Oyabu et al. performed *Amd1* and *Amd2* knockdown in primary murine myotubes that led to reductions in polyamine metabolism and atrophy [53]. They also highlight that dysfunctional polyamine metabolism is a common hallmark of muscle atrophy in mice [53]. We are the first to show, to our knowledge, that myofiber-specific *Smox* overexpression upregulates *Amd1/2* expression in myotubes. *Amd1/2* induction could help maintain cellular polyamine pools of spermine for its enzymatic conversion to spermidine, producing 5-methylthioadenoside precursors for methionine production via the methionine salvage pathway [53; 55].

Coinciding with higher *Smox* in both preconditioned plantaris myonuclei and mature myotubes was a reciprocal decrease in *Ddit4/Redd1*, a molecular repressor of mTORC1 and hypertrophy signaling [61; 62]. The repression of *Redd1* is sufficient to block atrophy in a variety of conditions [61; 90–92], providing evidence for its powerful role in muscle mass regulation. The suppression of REDD1 also accelerates loading-induced muscle hypertrophy [93]; a finding with clear implications for our own observations of accelerated hypertrophy due to exercise preconditioning. Furthermore, methionine metabolism associates to the expression of *Ddit4* expression during muscle wasting [62]. Changes to the methionine salvage pathway may be influenced by increased polyamine flux via *Smox* induction to affect *Ddit4* expression [55]. We therefore propose that exercise preconditioning potentially lowers the threshold for anabolic signaling. Rather than simply amplifying growth signals, previously trained muscle may have reduced REDD1-mediated restraint on mTORC1 alongside elevated SMOX and polyamine flux, together permitting a more rapid hypertrophic response. Further research is warranted to better understand the regulation of *Ddit4* as it pertains to SMOX and polyamine metabolism, especially in the context of exercise-induced muscle hypertrophy.

A limitation of the current study is that all of our analyses were performed in the basal (or resting) state in male mice, 48 hours after the last exercise bout. How preconditioned muscle responds sex-specifically to an acute exercise bout relative to naïve trained muscle was not evaluated. We speculate that many of the myonuclear epigenetic differences we observe with exercise preconditioning may have the most meaningful implications in the context of molecular responsiveness to an acute bout of exercise. Nevertheless, our findings collectively link capillarization, *Smox* induction, and *Ddit4/Redd1* suppression to myometaplasticity induced by prior exercise training. Our data are a roadmap for understanding how prior exercise modulates skeletal muscle adaptive potential, providing new potential targets for performance enhancement and therapeutic intervention.

## 4. MATERIALS AND METHODS

### 4.1 Animals and treatments

Animal procedures were approved by the University of Kentucky IACUC. Mice were housed in temperature- and humidity-controlled rooms, maintained on a 14:10-h light-dark cycle, and food and water were provided *ad libitum*. Male HSA^+/-^-GFP^+/-^ mice were generated by crossing homozygous human skeletal actin reverse tetracycline transactivator (HSA-rtTA) mice generated by our laboratory [17] with homozygous tetracycline response element histone 2B green fluorescent protein mice (TetO-H2B-GFP) obtained from Jackson Laboratory (005104). Mice were subjected to PoWeR as previously described [6; 8; 11; 94] and there were no differences in average daily running volume between groups, as previously reported [6]. Doxycycline (0.5mg/mL) in drinking water (2% sucrose) was given during the one week acclimation to the running wheel (unweighted “free wheel”) to label resident myonuclei with GFP. Wheels were locked in PoWeR training mice 48 hours before euthanasia, and all mice were fasted overnight prior to tissue collection. Mice were euthanized in the morning via a lethal intraperitoneal dosage of sodium pentobarbital followed by cervical dislocation. Plantaris muscles were harvested in the morning (before 11:00 AM), flash frozen in liquid nitrogen (LN_2_), and stored at -80°C until analyses. The plantaris muscle was chosen for multiomic analyses to assess the molecular regulation underlying enhanced hypertrophic responses after retraining compared to training naïve mouse muscle.

### 4.2 Bulk RNA isolation, sequencing, and analyses

Bulk RNA was isolated from plantaris muscle (5-10 mg) and C2C12 myotubes using cold TRIzol Reagent (Sigma-Aldrich, St. Louis, MO, USA). The isolated samples included biological replicates of DTRT (n=5) and CON (n=5) groups, and technical replicates of C21C12 myotubes transduced with a scrambled sequences myoAAV (n=3) and C2C12 myotubes transduced with *Smox* myoAAV (n=3). Samples were homogenized using zirconia beads and a Fisher Bead Mill (2 cycles of 15 seconds with 10 second stop in between). Following homogenization, RNA was isolated via phase separation by addition of chloroform then centrifugation. The aqueous phase was transferred to a new sterile tube and further processed on spin columns using the Direct-zol RNA MiniPrep kit (Zymo Research, Irvine, CA, USA; Cat #: R2050) [95; 96]. Concentration and purity were determined using a BioTek Take3 micro-volume microplate with a BioTek PowerWave XS microplate reader (BioTek Instruments Inc., Winooski, VT, USA).

Isolated RNA from all samples was sequenced by Novogene (Beijing, China) on an Illumina HiSeq using 150 bp paired-end sequencing, as previously performed [6; 96]. Raw FASTQ files were processed in Partek Flow and MyoAnalytics. Alignment was performed using STAR 2.7.8a, quantified to annotation model mm39, filtered for features at a cutoff of 10 counts, then normalization and statistical comparisons were performed with DESeq2. Genes with a false discovery rate (Benjamini–Hochberg method) adjusted *p*-value<0.05 were identified as differentially expressed genes (DEGs).

### 4.3 Mature C2C12 Myotube Differentiation and myoAAV Transduction

C2C12 myoblasts (2.6×10^3^ cells/mm) were plated on culture dishes and expanded in growth medium containing 10% fetal bovine serum (Gibco, 16000044) and 1% penicillin-streptomycin (Corning 30-002-CI) prepared in high glucose Dulbecco’s Modified Eagle Medium (DMEM) (Gibco, 11965-092). Media was replaced daily until ∼80% confluency, cells were differentiated using 2% horse serum (Gibco, 16050130), 1% penicillin-streptomycin (Corning), 5% HEPES (Gibco, 15630080), and 0.2% insulin-transferrin solution (Gibco, 41400-045) prepared in high glucose DMEM (Gibco) for 10 days. At day 10, cells were transduced at a multiplicity of infection (MOI) of 50,000 with myotropic adeno-associated viruses. A myoAAV (1C) delivering *Smox* cDNA and mCherry reporter (9.53×10^13^ GC/ml) or mCherry reporter only (1.93×10^14^ GC/ml) were designed and obtained through VectorBuilder (VectorBuilder Inc., Chicago, IL, USA). Myotubes were incubated with either experimental *Smox* myoAAV or control reporter-only myoAAV for 48 hours (days 10-12 of differentiation). After the induction period, myotubes were given only DM every 24 hours until myotubes were collected for analyses on day 14. Cells were used for RNA extraction, protein quantification, and myotube area normalized to myonuclear area as described by us previously [97].

For imaging, before seeding, cell culture plates were coated with a 2.5% gelatin with transglutaminase (10U/mL) and incubated overnight at 37°C to promote self-organization and alignment of C2C12 myotubes for fluorescence imaging [22] and immunolabeled for myosin heavy chain (MyHC; DSHB, MF20, 1:100) and stained for myonuclei (DAPI, 1:10000) in PBS with 2% BSA. Myotubes images from *Smox* cDNA (n=3) or scrambled sequences (n=3) transduced plates were acquired using a Zeiss microscope at constant exposures and intensity across technical replicates and images in the AF488 (MyHC), DAPI (myonuclei), and mCherry (myoAAV reporter) channels. Normalized myonuclear area was quantified from fluorescence images (GFP+ green channel) divided to the area of DAPI^+^ myonuclei (blue channel) using Fiji/Image J software.

For protein quantification, myotubes were harvested in 2X SDS sample buffer and assessed by Western blotting using anti-SMOX primary antibody (Cell Signaling Technology, 35842 1:1000). Myotube area statistical comparisons for SMOX protein abundance and relative myotube area were performed using one-tailed Welch’s t-tests with significance set a *p*<0.05, n=3 biological replicates per group, with values presented as mean ± SD.

### 4.4 Skeletal Muscle Capillarity (PECAM1) Immunohistochemistry

For immunohistochemistry (IHC) analyses, a distal portion of plantaris muscle was covered in Tissue-Tek optimal cutting temperature compound (OCT) (Sakura Finetek, Torrance, CA, USA) and pinned at resting length to a cork covered in aluminum foil. Muscles were frozen in liquid nitrogen-cooled isopentane and stored at −80°C. At the time of sectioning, muscles were oriented upright in tissue OCT, which was then frozen using freeze spray in order to secure the muscle sample in place. Plantaris muscle sections were cut on a cryostat at −24°C. Frozen muscle sections (7μm) were air-dried for >30 minutes and stored at −20°C.

The PECAM1 (Cat#: 2H8-supernatant) monoclonal antibody developed and deposited to the Developmental Studies Hybridoma Bank by Bogen, S.A., created by the NICHD of the NIH and maintained at The University of Iowa, Department of Biology, Iowa City, IA 52242 and used for determination of plantaris muscle capillarity. For PECAM1, plantaris muscle sections were removed from -20°C and air dried for 15 mins then fixed in cold acetone (-20°C) for 10 mins followed by 2×5 min washes in PBS. Sections were then blocked for 30 min in 5% normal horse serum (NHS) in PBS. Primary antibodies for PECAM1 (1:5) and dystrophin (1:250, PA1-37587, Invitrogen, Carlsbad, CA, USA) in PBS with 5% NHS were completed for 1hr at room temperature then overnight at 4°C. The following day sections were washed 3×5 min in PBS and then incubated in secondary antibodies against PECAM1 (1:500, AF594, goat anti-Armenian hamster IgG, Invitrogen) and dystrophin (1:200, AF350, donkey anti-rabbit IgG, Invitrogen) for 1hr at room temperature. Following secondary incubation, sections were washed 3×5 mins with PBS and mounted with VectaShield fluorescent mounting media (Vector). Images were captured at 20x total magnification at room temperature with a ZEISS upright microscope (Axio Imager M2, Oberkochen Germany). For analysis of capillarization, each sample had multiple ROIs (225×225µm) captured (2-3 per sample) for analysis of capillary density (capillaries per mm^2^). Image analysis was performed by the same investigator who was blinded to sample condition throughout the analysis using Zeiss ZEN microscopy software. Limitations in sample availability in the CON group led to a reduction in its sample size (n=3) for analysis with no sample exclusion in the DTRT group (n=5). Unpaired t-test with test for normality and significance value set at *p*<0.05 were used to determine differences in capillarization between preconditioned and naïve trained plantaris muscle.

### 4.5 Single Myonuclei Isolation

Isolation of single myonuclei was completed via fluorescent activated nuclear sorting (FANS) with the MACSQuant Tyto Cell Sorter (Miltenyi Biotec, Bergisch Gladbach, Germany) as we have described previously [98]. Plantaris muscle samples were used to isolate myonuclei for assessment of DNA methylation and single myonucleus RNA-sequencing. For DNA methylation, plantaris muscle (∼10 mg) was placed in a small glass beaker with a sucrose-based buffer to mimic physiological conditions (5mM PIPES, 85mM KCl, 1mM CaCl_2_, 5% sucrose, 2X Halt Protease inhibitors, and 0.25% NP-40), pulsed with propidium iodine (PI) to fluorescently label DNA for viability, then minced into a slurry with scissors. The nuclear suspension was then transferred to a manual glass Dounce for manual homogenization, then strained through a 20µm MACSQuant pre-separation filter directly into MACSQuant Tyto regular-speed sorting cartridge (Cat #: 130-104-791). FANS gating was established to removed debris and identify nuclei positive for both GFP^+^ (intrinsic/resident myonuclear label) and PI^+^, which was sorted into ATL buffer with proteinase K (Qiagen) for downstream DNA extraction and DNA methylation analysis. For single myonucleus RNA sequencing experiments, 2-3 plantaris muscle samples were pooled (∼15-20mg total per pooled sample, n=2 pooled samples for each condition [DTRT and CON], n=2-3 mouse plantaris per pool) in homogenization buffer (500µL HEPES [1M], 3mL KCl [1M], 250µL spermidine [100mM], 750µL spermine tetrahydrochloride [10mM], 10mL EDTA [10mM], 250µL EGTA [100mM], 2.5mL MgCl [100mM], 5.13g sucrose). Samples were minced into a slurry with scissors and manually homogenized with a plastic pestle in a low-bind 1.5mL tube in a sucrose buffer mimicking physiological conditions with RNase inhibitors. The nuclear suspension was then strained through a 20-µm MACSQuant pre-separation filter directly into MACSQuant TYTO high-speed sorting cartridge. Myonuclei were sorted into 1X PBS, 1% BSA, with RNase inhibitors.

### 4.6 Myonuclear Reduced Representation Bisulfite Sequencing (RRBS)

DNA was extracted from the isolated myonuclei of the plantaris using a QIAamp® DNA Micro Kit (Germantown, MD, USA; Cat #: 56304). Briefly, sorted myonuclei in buffer ATL and proteinase K were incubated for ≥4 hours at 56°C. DNA binding to the column was conducted using 1 µg of carrier RNA, and washes and centrifugations were carried out according to the manufacturer’s instructions. DNA was eluted in 12 µl of nuclease-free H_2_O, quality checked on the Agilent Tapestation with the genomic DNA (gDNA) screen tape to verify presence of large DNA fragment size (generally <30kb) and placed in -20°C until later analyses. RRBS was performed by Zymo Research using ≥10 ng of gDNA. Some samples did not reach the minimum gDNA mass requirement for RRBS and could not be included in downstream analysis (DTRT n=1; CON n=1). RRBS data were processed using methylKit [99] with a minimum cut-off of 10x coverage per CpG site in each sample, as previously described [6; 96]. Promoter-associated CpGs were defined as within ±10% of the corresponding gene length surrounding the transcriptional start site (TSS) [6]. Differentially methylated sites were determined using logistic regression with a chi-square test and sliding linear model (SLIM)-adjusted *q*-values are reported, with *q*<0.05 considered significant unless stated otherwise. Differentially methylated CpGs at promoter regions (adj. *p*<0.05) were uploaded into ShinyGO as gene symbols (http://bioinformatics.sdstate.edu/go/) [100] and over-representation analyses for KEGG pathways were performed using a cutoff of FDR<0.05. Figures were extracted from ShinyGO. Results are presented at the gene level, and site-specific CpG data are provided in the Supporting Information (Supplemental Table S1).

### 4.7 10X Single Myonucleus RNA Sequencing Library Preparation and Analysis

GFP^+^/PI^+^ myonuclei were isolated via FANS on our MACSQuant Tyto sorter. Due to the low nuclear yield from FANS, we combined the maximum volume of nuclei suspension (up to 36.6 µl) with 28.4 µl of reverse transcription master mix (MM). The myonuclei + MM solution was loaded into the 10X Chromium system using the Single Cell 3’ Reagent Kit v4 according to the manufacturer’s protocol. Following library construction, libraries were sequenced on the Illumina Nova NextSeq X Plus System by Novogene to 200 million reads per sample. Raw FASTQs were imported into Cell Ranger 9.0 to generate feature-barcode matrices. Raw-feature h5 files were then processed in a Python environment using CellBender [101] to remove-background for ambient RNA correction, using sample-specific estimates of expected nuclei and total droplets. Processed files were imported into Seurat v5 [102] for downstream processing. Genes detected in <3 nuclei and nuclei with <100 detected genes, <200 UMIs, or >5% mitochondrial reads were removed. Datasets were normalized using SCTransform(), followed by principal component analysis, nearest-neighbor graph construction, clustering, and UMAP dimensionality reduction. Doublets were identified and removed using scDblFinder [103]. Singlet datasets were then integrated with SCT normalization using the top 3000 variable features, followed by PCA-guided dimensionality reduction, UMAP embedding, and Leiden clustering [104], with the final clustering resolution selected based on modularity and silhouette metrics. For downstream differential expression analysis, normalized RNA expressions were compared between exercise preconditioned and training naïve myonuclei within annotated populations using the Wilcoxon rank-sum test. Gene set enrichment analysis was performed using clusterProfiler for KEGG and Gene Ontology Biological Process pathways (see Supplemental Table S2). Genes were ranked using the following: Rank Score = sign(log2FC)*-log10(p-value), and pathways with adjusted p <0.05 were considered significantly enriched.

### 4.8 Binding and Expression Target Analysis (BETA)

Pathway analysis was performed using the enrichR R-package. Up- and down target genes resulting from BETA integration of DNA methylation and RNA-seq datasets were included in independent up and down gene sets as previously described [20; 21; 98]. Briefly, the underlying algorithm of BETA takes into consideration the distance of the regulatory element relative to the TSS by modeling the effect of regulation using a natural log function, termed the regulatory potential, as described by Tang et al. [105]. CpG sites within the defined promoter-associated regions were converted into ‘methylation peaks’ as a surrogate for transcription factor binding peaks, with the significance scores calculated as the -log_10_(*q*-value). To determine whether methylation has regulatory potential, a non-parametric statistical test contrasted regulatory potentials for differentially methylated CpGs upstream or downstream of the TSS with differentially expressed genes in the experiment (in this instance, DTRT vs. CON defines the differential expression). Alpha (*a*) for the BETA integration analysis was set at *p*<0.05, and individual comparisons were reported using rank product analysis. It is worth noting that the genomic location information for CpGs from the array was used for the BETA input and given the nature of the BETA algorithm is based on regulatory potential, sometimes regulation of genes that differed from the genes that were classified in the array was identified, which is purely based on distance to the TSS [20; 46].

## Supplemental Table Legends

**Table S1.** Methylation at CpGs sites in promoter regions in exercise preconditioned myonuclei. Results describe the CpG coordinates (coords), seq name, CG start and end, and *p-* and q-values for the comparison between DTRT and CON, methylation difference, individual percent methylation for each CpG in each study sample, and gene symbol.

**Table S2.** Single myonucleus RNA-sequencing data sets for each myonuclear cluster (coarse, spindles, Type IIa, Type IIb, Type IIx, NMJ, and MTJ) in exercise preconditioned myonuclei relative to training naïve myonuclei. Sequencing results include gene symbol, log_2_FC, *p*-value, and adj. *p*-value. KEGG and GOBP pathways enrichment (adj. *p*<0.05) results include pathway ID, description, set size, enrichment, normalized enrichment score (NES), *p*-value, adj. *p-*value, q-value, rank, leading edge, core enrichment, and core enrichment gene symbols.

**Table S3.** Transcriptomic profile of *Smox* myoAAV induced C2C12 mature myotubes relative to controls. Sequencing results include gene ID, gene symbol, p-value, gene expression by sample, and average group gene expression.

**Table S4.** Complete lists of unique and shared DEGs between the *in vitro Smox* overexpression transcriptome and *in vivo* myonuclear profiles with preconditioning. Shared (C2C12+smnRNA-seq) and unique (C2C12 or smnRNA-seq only) expression profiles for differentially expressed genes compared to respective controls are presented.

## Author Contributions

**T.L.C.**, **K.A.M.** and **Y.W.** conceived the study. **T.L.C., A.R.C., R.G.J., P.J.K., F.M., N.S., A.P.**, **Z.B.M, F.v.W, C.M.D, and K.A.M.** performed experiments and/or analysis. **A.R.C., K.P., D.A.E., A.L.S.,** and **C.E.N**. contributed to the *in vitro* experiments. **K.A.M, Y.W., D.A.E., F.v.W,** and **C.E.N**. provided resources. **T.L.C**. and **K.A.M**. wrote the manuscript draft with input from **R.G.J.** and **Y.W.** All authors reviewed and approved the manuscript.

## Ethics Statement

All animal procedures were approved by the Institutional Animal Care and Use Committee of the University of Kentucky.

## Consent

The authors have nothing to report.

## Data Availability Statement

The data that support the findings of this study are available in the supplementary material of this article. RRBS, bulk RNA-seq, and smnRNA-seq data sets will be deposited in the Gene Expression Omnibus (GEO) upon publication.

